# Using nonthermal plasma treatment to improve quality and durability of hydrophilic coatings on hydrophobic polymer surfaces

**DOI:** 10.1101/868885

**Authors:** Greg D. Learn, Emerson J. Lai, Horst A. von Recum

**Author notes:** Corresponding Author, 10900 Euclid Avenue, Cleveland, OH 44106.

## Abstract

Low surface energy substrates, which include many polymers in medicine/industry, present challenges toward achieving uniform, adherent, durable coatings, thus limiting intended coating function. Examples include hydrophobic polymers such as polypropylene, polyethylene, polytetrafluoroethylene, and polydimethylsiloxane. These inert materials are used in various biomedical implants due to favorable bulk properties despite perhaps unfavorable surface properties. The capability to coat such materials holds great value as the surface heavily influences biological response and implant function *in vivo*. Likewise, paint/ink coatings are often necessary on these same plastics, as their final appearance can be critical for automotive, packaging, and consumer products. Substrate exposure to nonthermal plasma was explored here as a means to improve quality of coatings, specifically cyclodextrin-based polyurethanes previously explored for biomedical applications such as controlled drug delivery and anti-biofouling, upon otherwise incompatible polypropylene substrates. Plasma treatment was found to increase wettability and oxygen content on substrate surfaces. These plasma-induced surface alterations were associated with enhanced coating uniformity, and improved coating/substrate adherence – determined to derive partly from interfacial covalent bond formation. Findings demonstrate the utility of plasma-based surface activation as a strategy to improve coating quality on polymeric substrates, and reveal insights regarding mechanisms by which plasma improves polymer coating adherence.

## 1. Introduction

The surface plays a critical role in the success and performance of many materials, including surgically implanted devices. Events such as protein adsorption, cell or bacterial attachment, biofilm formation, blood coagulation, tissue adhesion, foreign body response, and corrosion can all take place at the interface between an implant and the human body. Therefore, an implant material should not be selected for a particular medical application on the basis of its bulk properties alone. A large body of research has been directed toward modifying implant material surfaces to achieve desirable host-material interactions without compromising bulk properties. One of the most common surface modification approaches used on implant materials is the application of coatings, a strategy which is advantageous in that bulk material properties are expected to be retained while the product obtains desirable surface properties consistent with the coating material. On medical devices, these coatings may serve many purposes including reduction of non-specific protein adsorption or bacterial attachment, sustained drug release, enhancement of attachment of certain host cells, and prevention of corrosion.

A large number of biomedical implants are composed of polymers that possess low surface energy. Such polymeric materials include: 1) polypropylene (PP), present in surgical meshes and sutures, 2) polyethylene (PE), found in total joint replacements, 3) polydimethylsiloxane (PDMS), utilized in plastic surgery, and 4) polytetrafluoroethylene (PTFE), used for catheters and arterial grafts. The low surface energy of these materials is disadvantageous for several reasons: (i) it promotes adsorption and denaturation of proteins on the bare surface, along with subsequent inflammatory responses, as it is favorable for proteins to displace water at the hydrophobic surface and change conformation to allow core nonpolar amino acids to associate with the material^1^, and (ii) it decreases the receptiveness of these materials toward coatings^2^, which could otherwise allow for improved host response and device function. In general, the surface energy of a substrate should exceed that of a coating to achieve reasonable spreading and adhesion. If not, coatings applied to low surface energy substrates suffer from lack of uniformity, adherence, and durability, limiting the intended function of the coating over the lifetime of the implant.

An attractive solution to promote uniformity and adherence of coatings on such difficult materials is treatment of the substrate with nonthermal plasma^3,4^, a state of matter created when a gas becomes ionized through application of sufficient energy in the form of electromagnetic fields. Plasma is composed of a mixture of positive and negative ions, neutral atoms/molecules, radicals, free electrons, and photons. Interaction of plasma with a polymer results in surface cleaning/etching, scission or rearrangement of bonds, and the introduction of new functional groups as determined by the composition of the carrier gas. These plasma-induced changes can contribute to improved substrate receptiveness to coatings^5^, and also directly alter biological responses at the substrate surface^6^. Furthermore, plasma treatments are amenable for substrates with complex geometries, circumvent the need for added hazardous chemicals as adhesion promoters (e.g. solvents), and can achieve desirable surface changes whilst minimizing impact on the bulk properties of the substrate material^6^.

Our group has previously developed a polysaccharide-based material as a coating that can serve as a vehicle for sustained drug delivery^7–10^ and also mitigate events of biofouling such as protein adsorption and bacterial attachment^11^. This unique polymer is synthesized from subunits of cyclodextrin, an excipient found in some pharmaceutical formulations. Coatings of polymerized cyclodextrin (pCD) have previously been applied to polyester surgical fabrics, silicone catheters, and metallic orthopedic screws^7,8,12–14^. However, these previous coatings, while successful, were difficult to control in terms of uniformity and adherence^15^. Furthermore, given that many inert polymers are used as implant materials in medicine, and the need for functional and stable biomaterial coatings, it would be advantageous to fundamentally explore the application of pCD coatings upon these more difficult substrates, so as to maximize coating uniformity and adherence. The objective of this work, therefore, is to investigate the effects of nonthermal plasma activation of PP substrates on the quality of pCD coatings. PP is chosen as a model substrate material given its inherently low surface energy, and its use in many medical products, such as surgical sutures and meshes, implants for which pCD coatings may be useful for mitigating inflammatory, infective, and adhesive complications.

An overview of this work is presented in **Figure 1**. The hypothesis of this study is that nonthermal plasma treatment enhances the uniformity and adherence of pCD coatings on PP substrates. To test this hypothesis, the time-dependent effects of nonthermal plasma exposure on PP surface characteristics were first evaluated using contact angle goniometry and X-ray photoelectron spectroscopy (XPS). Next, the effects of substrate plasma activation on pCD coating uniformity and adherence were investigated. Uniformity was assessed both macroscopically and through the use of scanning electron microscopy (SEM). Adherence was evaluated by way of lap-shear testing according to ASTM standard. Finally, to corroborate findings regarding effects of substrate plasma exposure on coating adherence, XPS was performed to evaluate formation of covalent bonds between functional groups at the coating-substrate interface.

**Figure 1:**
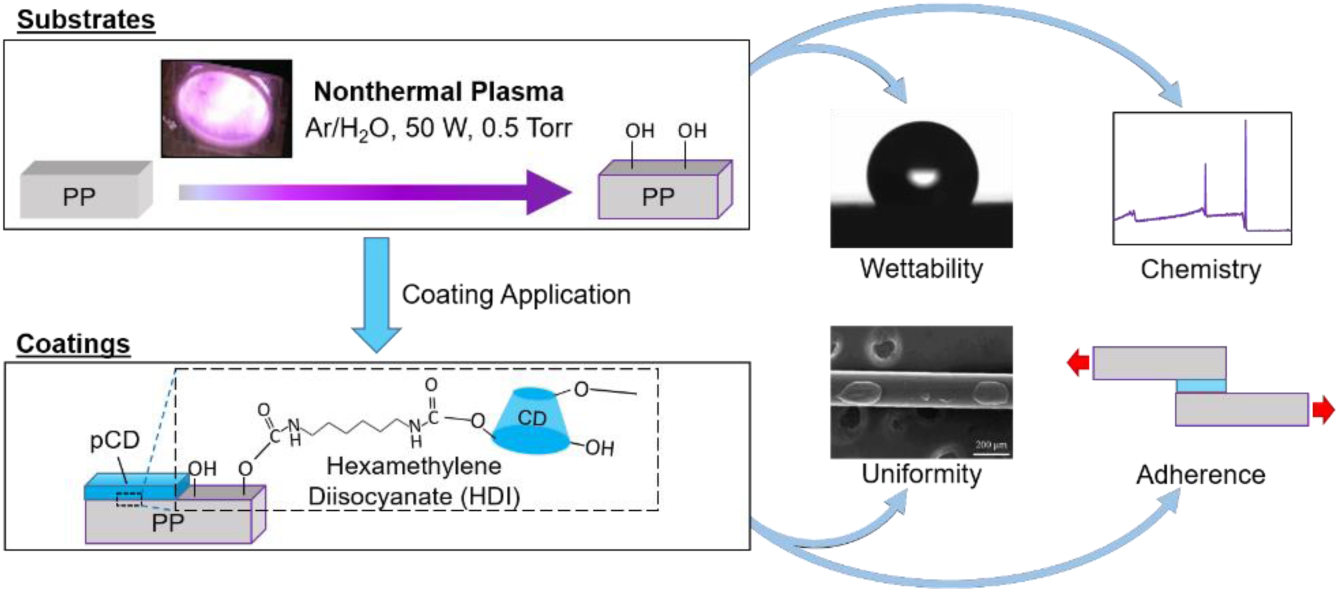
Study overview. Effects of nonthermal plasma on PP substrates were investigated in terms of wettability and surface chemistry, and effects on pCD coatings were explored in terms of uniformity and adherence.

## 2. Methods

### 2.1. Materials

Soluble, lightly epichlorohydrin-crosslinked β-CD polymer precursor (bCD) was purchased from CycloLab R&D (#CY-2009, batch CYL-4160, MW ∼116 kDa; Budapest, Hungary). HDI crosslinker (#52649) and 2-(trifluoromethyl)phenyl isocyanate (2-TPI) (#159379) were purchased from Sigma Aldrich. N,N-dimethylformamide (DMF) solvent (#D119-4) was purchased from Fisher Scientific. PP 24-well plates (#1185U58) and lids (#1185U62) for macroscopic coating observation were purchased from Thomas Scientific. PP sheet stock (#8742K133) for XPS and contact angle goniometry, and bar stock (#8782K11) for lap-shear testing were purchased from McMaster-Carr. PP 4-0 Prolene blue suture (#8592G) was purchased from eSutures.

### 2.2. Plasma cleaning and activation of PP substrate surfaces

In order to systematically examine effects of plasma treatment on pCD coatings and PP substrates, PP materials were placed in a 4” (10 cm) diameter x 8” (20 cm) length quartz reaction chamber of a Branson/IPC Model #1005-248 Gas Plasma Cleaner and treated with low-pressure nonthermal plasma (500 mTorr, 50 W, 13.56 MHz) using an inlet gas mixture of argon bubbled through water (Ar/H_2_O). Effects of plasma treatment on PP substrate wettability and surface chemistry were investigated using treatments of durations from 1-20 min performed within 6 h or 12 h of contact angle measurement or XPS analysis, respectively. The impact of plasma treatment on pCD coating uniformity and adherence was assessed using a fixed substrate treatment duration of 10min within 1 h of pCD coating application. Non-treated (0min) PP samples without any known prior exposure to plasma or ultraviolet light were included as controls in all experiments.

### 2.3. Effects of plasma treatment on PP substrates

#### 2.3.1. Contact angle goniometry

For polar pCD coatings to spread uniformly on nonpolar PP substrates, the surface must be rendered more wettable, therefore the effect of plasma treatment on wettability of PP was examined using contact angle goniometry. PP sheet stock was cut to dimensions of ∼1.5 × 1.5 in^2^ (38 × 38 mm^2^), gently sanded to expose fresh surface using a graded series of SiC sandpaper (1200, 2500, and 5000 grit), and rinsed thoroughly with deionized water prior to performing plasma treatments (0, 1, 2.5, 5, 10, and 20 min). After sanding and/or plasma treatments, care was taken to ensure that faces to be analyzed were not inadvertently exposed directly to any liquid or solid materials before analysis. PP surfaces were then evaluated for wettability by static contact angle measurement using a KSV Instruments CAM 200 Optical Contact Angle Meter. Deionized water droplets (n=9-14 unique droplets per sample) of 8 µL volume were dispensed onto each PP surface and allowed to equilibrate for 30 s prior to photographing and measurement. The measurement for each droplet reflects the average of the angles on the left and right sides. Measurements were performed using KSV CAM 2008 software. Results shown represent findings from one experiment, with the same trends having also been observed in 2 similar independent experiments.

#### 2.3.2. X-ray Photoelectron Spectroscopy

Effects of plasma treatment on PP surface chemistry were studied using XPS to better understand the mechanistic basis by which plasma impacts spreading and adhesion of pCD coatings onto PP substrates. PP sheet stock was cut to dimensions of ∼7.5 × 7.5 mm^2^, gently sanded to expose fresh surface using a graded series of SiC sandpaper, and rinsed thoroughly with deionized water prior to performing plasma treatments (0, 1, 2.5, 5, 10, and 20 min). After sanding and/or plasma treatments, care was taken to ensure that faces to be analyzed were not inadvertently exposed directly to any liquid or solid materials before analysis. PP surfaces were then analyzed for elemental content using a PHI Versaprobe 5000 Scanning X-Ray Photoelectron Spectrometer equipped with Al Kα source (hν = 1486.6 eV). Scans were acquired on a total of n = 3-4 samples per plasma treatment duration across 2 pooled experiments, with 2 unique scan locations averaged per sample. Survey scans were collected using a 200 µm spot size, 45 W power, 15 kV acceleration voltage, 117.40 eV pass energy, 0.40 eV step size, 25 ms/step, 8 cycles, 44.7° take-off angle, and 0-1100 eV range. The C1s peak was auto-shifted to 284.8 eV, and the ratios of the elements carbon, nitrogen, and oxygen were analyzed. The areas of peaks were taken with background set using a Shirley function from 280-292 eV for C1s, 396-404 eV for N1s, and 526-538 eV for O1s. Auger peaks were not used for analysis. After survey scans, high-resolution scans were collected using a 100 µm spot size, 25.2 W power, 15 kV acceleration voltage, 23.50 eV pass energy, 0.20 eV step size, 50 ms/step, 16 cycles, 44.7° take-off angle, and 278-298 eV range for C1s, or 523-543 eV range for O1s. The vertical sampling depth ζ (from which 95% of signal originates) for take-off angle θ = 44.7° and reported inelastic mean free path of λ = 3.5 nm at a photoelectron kinetic energy of 1 keV for PP surfaces^16^, is determined to be ∼7.4 nm based on the relation^17^ ζ = 3λcos(θ). Analysis was performed using MultiPak software version 9.8.0.19 (Physical Electronics, Inc.). For interpretation of high-resolution scans, C-C, C-O, C=O, and O-C=O peak positions^18–22^ were constrained at 284.8±0.1, 286.3±0.1, 288±0.2, and 289±0.2 eV, respectively, and full widths at half maximum (FWHM) were constrained to be within 10% that of the C-C peak FWHM value.

### 2.4. pCD synthesis and coating onto surfaces

pCD coatings were synthesized in three steps using HDI as a crosslinker for bCD. First, one gram of bCD was weighed and placed in a PP tube, then 3 mL DMF was added to dissolve it. Second, HDI was added so as to initiate crosslinking, and mixtures were thoroughly vortexed. Third, pre-polymer mixtures were cast as coatings either: (i) into wells of PP multiwell plates for production of coated well surfaces for visualization of coatings, (ii) onto PP sutures for visualization of coatings under SEM, or (iii) onto flat PP bar stock pieces (newly abraded using SiC sandpaper to expose fresh surface) for preparing coated specimens for lap-shear testing. For coated well surfaces the amount of HDI crosslinker was varied between extremes of 80 µL and 640 µL per gram bCD, to achieve approximate crosslink molar ratios (HDI per glucose residue) of 0.08 and 0.64, respectively, chosen to span a range from the minimum limit for gelation up to brittle materials. The amount of HDI used in all other experiments was 320 µL per gram bCD to achieve a crosslink ratio of 0.32, chosen as an intermediate value. For coated well surfaces, the volume of pre-polymer mixture added was 140 µL/well for 24-well plates, then all plates were agitated gently to help promote complete coverage. Cast pre-polymer mixtures were kept covered with Parafilm and allowed to cure for at least 2 days at ambient temperature and pressure. Cured pCD coatings were rinsed several times to terminate crosslinking, and stored immersed in deionized water to keep samples hydrated before use.

### 2.5. Effects of plasma treatment on pCD coatings

#### 2.5.1. Scanning Electron Microscopy

SEM was performed to study pCD coating uniformity on PP substrates with or without prior plasma surface activation. Prolene sutures were removed from sterile packaging, cut into 1” long pieces, and either treated with nonthermal plasma for 10 min or left untreated. Sutures were then coated with pCD by dipping in pre-polymer mixture, then covered and allowed to cure for 2 days at ambient temperature and pressure. pCD-coated sutures were then gently adhered to a stub using carbon tape, and sputter-coated with 5nm of palladium under vacuum. Sutures were characterized using a JSM-6510 series JEOL scanning electron microscope. Images were taken at 50x magnification and an excitation voltage of 25kV.

#### 2.5.2. Lap-shear testing of pCD coatings on PP substrates

To examine the impact of plasma surface activation on the adhesion of PP substrates to pCD coatings, lap-shear testing was performed. Lap-shear testing and specimen preparation were performed in accordance with the ASTM D3163-01(2014) standard^23^. pCD pre-polymer mixtures (150 µL) were cast between two rectangular strips of PP sheet substrate (4” × 1” × 1/8”) on a 1” × 1” overlap region, and allowed to cure for 6 days at ambient temperature and pressure. Paired PP strips were either left untreated, or treated with nonthermal plasma for 10 min, prior to pCD application. Overlap regions were securely held together over the curing period with the use of paired medium-size binder clips and excess pre-polymer that spilled out of the joint was wiped off with a KimWipe. Lap joints (n = 5/group) were tested in tension until failure at a rate of 1.3 mm/min (0.05”/min) and a sampling rate of 72 Hz on an Instru-Met renewed load frame operated with Testworks 4 software and hand-tightened vice grips capable of being horizontally offset so as to minimize peel effects. Load was monitored using a 100 lbf tension load cell. The maximum load attained during the test was divided by the overlap area and recorded as the lap-shear strength. The work to failure was divided by the overlap area and recorded as the lap-shear toughness. Grips were maintained with 2.5” between the grip edge and the bonded region (0.5” specimen/grip overlap). Results shown represent findings from one experiment, with the same trend having also been observed in one similar independent experiment.

### 2.6. Effects of plasma treatment on covalent bonding at coating-substrate interfaces

In order to better understand the mechanisms behind the effects of plasma treatment on pCD coating adhesion to PP substrates, we sought to determine whether covalent bonding occurs between pCD coatings and the plasma-treated PP surface. However, to do this an isocyanate compound was needed that could be uniquely identified at the activated PP surface. This was done using XPS to detect fluorine as a unique marker of the isocyanate compound 2-TPI after exposure of this compound to untreated and plasma-treated PP surfaces. PP sheet stock was first cut to dimensions of ∼7.5 × 7.5 mm^2^, gently sanded to expose fresh surface using a graded series of SiC sandpaper, and rinsed thoroughly with deionized water prior to performing plasma treatments (0 versus 10 min). Sample surfaces were then directly exposed to 2-TPI overnight, followed by two sequential rinses with copious amounts of toluene, acetone, isopropanol, then deionized water. Finally, a stream of dry nitrogen gas was used to remove excess water from sample surfaces. Untreated and plasma treated PP surfaces that were not exposed to isocyanate were prepared as controls as well. After sanding, plasma, and isocyanate exposure, care was taken to ensure that faces to be analyzed were not inadvertently touched by any solid surface before or during analysis. PP surfaces were then analyzed for elemental content using XPS as above. XPS was performed within 48h of plasma treatment. Survey scans were acquired on a total of 2 samples per plasma treatment duration, with 2 unique scan locations per sample. The ratios of carbon, nitrogen, oxygen, and fluorine were analyzed using Multipak software. The areas of peaks were taken with background set using a Shirley function as before for C1s, N1s, and O1s, and from 675-695 eV for F1s. Auger peaks were not used for analysis. Increased fluorine content was attributed to covalent urethane bond formation between hydroxyl groups on the PP surface and the isocyanate group on 2-(trifluoromethyl)phenyl isocyanate.

### 2.7. Statistical analysis

All data is presented as mean ± standard deviation. Statistical analysis tests were carried out in Microsoft Excel 2016. For all comparisons, two-sample two-tailed Student’s t-tests with unequal variance were used. Statistical significance was set at p < α = 0.05.

## 3. Results

### 3.1. Effects of plasma treatment on PP substrate wettability and surface composition

First, we sought to evaluate the effects of plasma on the substrate surface, particularly in terms of wettability and composition. Wettability was studied through contact angle goniometry using the sessile drop method. Plasma treatment for any length of time was found to increase the wettability of PP substrates (p < 0.001) (**Table 1**), with longer treatments correlating with decreased static water contact angles. Without plasma treatment, water contact angles on PP averaged >120°, but after 20 min of plasma exposure, this value decreased to <60°.

PP substrate surface composition was studied using XPS. Quantitative analysis of survey scan spectra (**Fig 2**) indicated that plasma exposure for any duration enhanced the amount of oxygen on PP surfaces (p < 0.001), and decreased the level of carbon (p < 0.001), with no significant impact on surface nitrogen content (p > 0.1) until 10 or 20 min (p < 0.014) of treatment (**Table 1**). Longer treatments were associated with increased oxygen, slightly increased nitrogen, and decreased carbon concentrations. High-resolution C1s spectra were nearly symmetric for the untreated PP surface, but demonstrated an increasing skew to the left as plasma exposure time increased (**Fig 3**), suggesting increasing abundance of oxygen-containing functionalities including C-O, C=O, and O-C=O. Deconvolutions of high-resolution C1s scans suggested that most engrafted oxygen was incorporated as hydroxyl groups, with lesser amounts as ketones/aldehydes and carboxyls/esters, for all tested plasma durations. This finding is in agreement with previous reports^18^. However, the abundance of hydroxyl groups seemed to level off after 10 min of treatment, as existing hydroxyls were likely oxidized further to species such as ketones, carboxylic acids, or ultimately even carbonates^19,20,22^. For this reason, a treatment duration of 10 min was utilized from this point in all subsequent experiments. Engraftment of oxygen-containing groups likely explains the enhanced surface wettability of PP following plasma treatment.

**Table 1:**
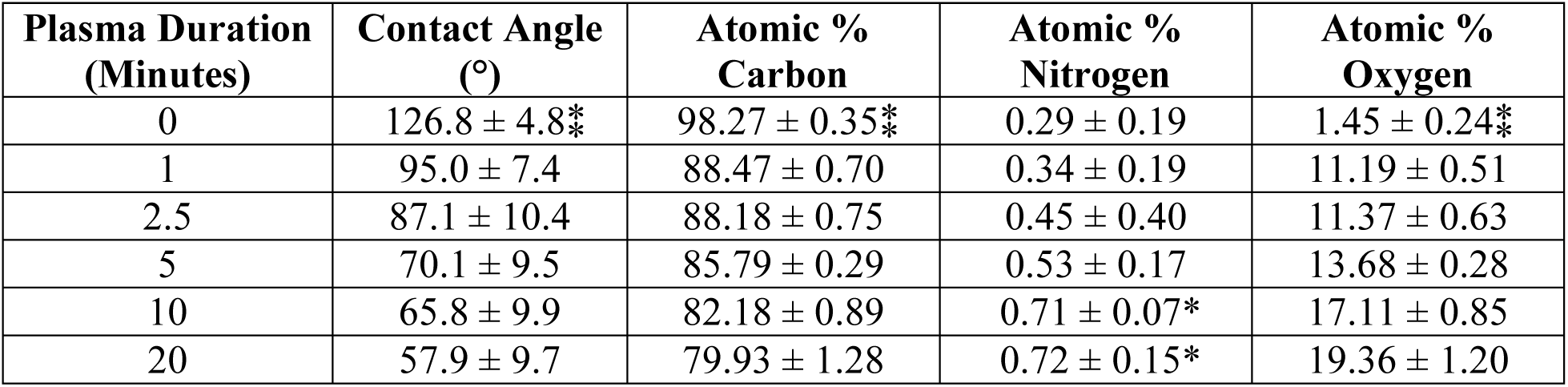
Effect of plasma treatment duration on PP wettability (n = 9-14 droplets per time point), and surface chemistry (n = 3-4 samples per time point). ⁑Significant difference to all subsequent time points. *Significant difference to 0 min.

| Plasma Duration (Minutes) | Contact Angle (°) | Atomic % Carbon | Atomic % Nitrogen | Atomic % Oxygen |
| --- | --- | --- | --- | --- |
| 0 | 126.8 ± 4.8* | 98.27 ± 0.35* | 0.29 ± 0.19 | 1.45 ± 0.24* |
| 1 | 95.0 ± 7.4 | 88.47 ± 0.70 | 0.34 ± 0.19 | 11.19 ± 0.51 |
| 2.5 | 87.1 ± 10.4 | 88.18 ± 0.75 | 0.45 ± 0.40 | 11.37 ± 0.63 |
| 5 | 70.1 ± 9.5 | 85.79 ± 0.29 | 0.53 ± 0.17 | 13.68 ± 0.28 |
| 10 | 65.8 ± 9.9 | 82.18 ± 0.89 | 0.71 ± 0.07* | 17.11 ± 0.85 |
| 20 | 57.9 ± 9.7 | 79.93 ± 1.28 | 0.72 ± 0.15* | 19.36 ± 1.20 |

**Figure 2:**
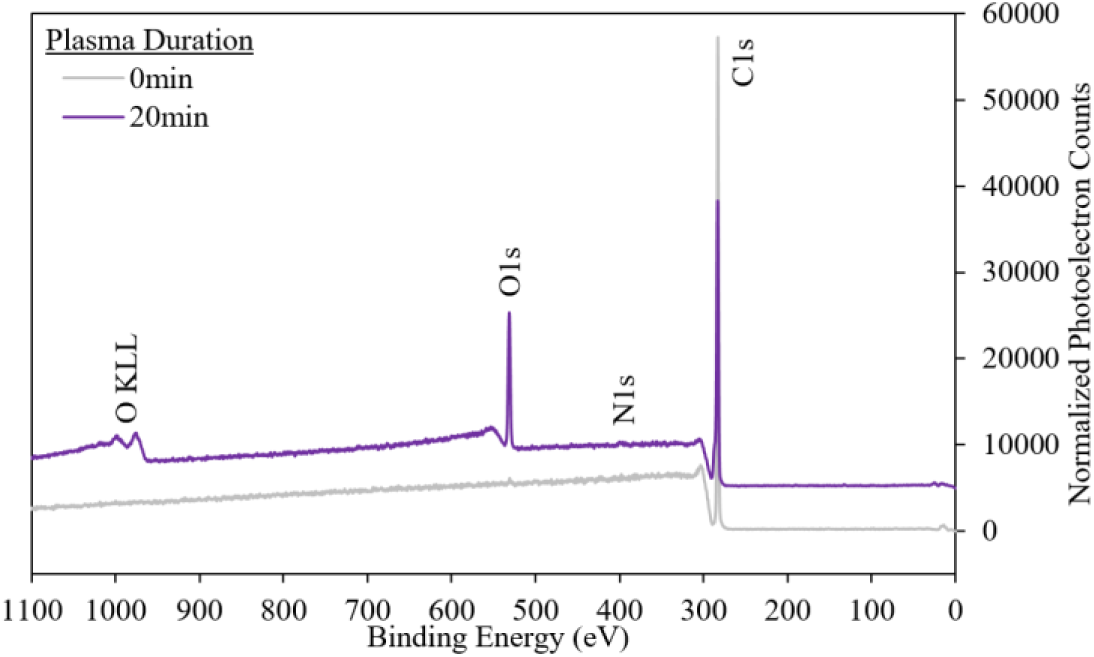
Representative XPS survey scans (rescaled to normalize areas under curves) of the 0 and 20 min time points demonstrate increasing height of the O1s peak with plasma treatment. KLL peaks indicate atomic relaxation via the Auger effect, and were not used for analysis. Spectra are intentionally offset along the ordinate.

**Figure 3:**
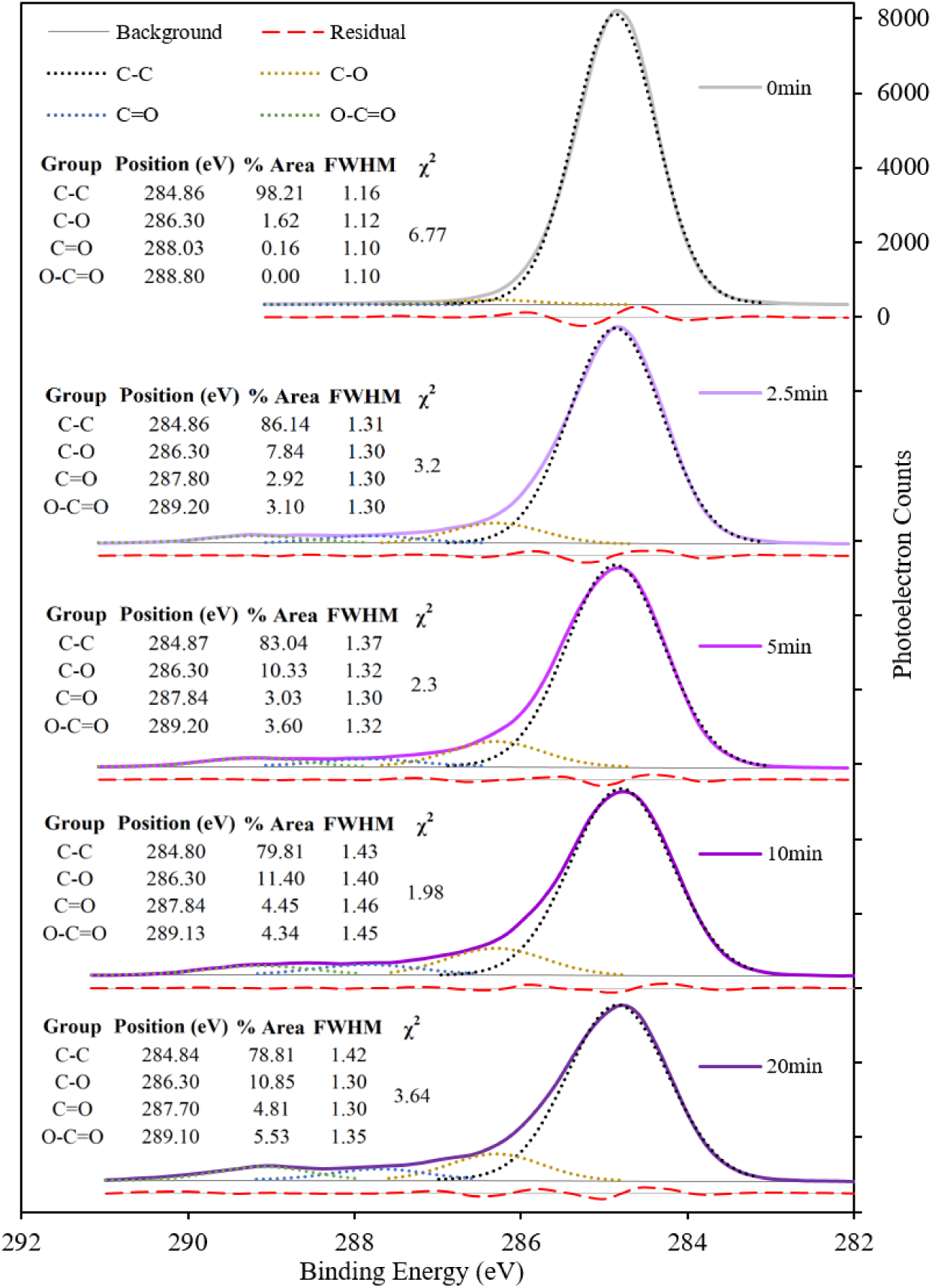
Representative XPS high-resolution scans demonstrate increasing abundance of oxygen-containing functional groups with increasing plasma treatment time. Spectra are intentionally offset along the ordinate.

### 3.2. Effects of plasma treatment on pCD coating uniformity and adherence

Having characterized the effects of plasma treatments on PP substrates, we next aimed to evaluate the effects of the plasma on pCD coating uniformity and adherence. Coating uniformity was assessed based on morphologic appearance, and using SEM. pCD coatings uniformly and stably covered the surfaces of PP substrates but only after plasma treatment. This was observed both macroscopically on PP 24-well plate surfaces (**Fig 4**), and microscopically on PP suture surfaces (**Fig 5**). Without prior substrate plasma treatment, coatings tended to bead up at well edges, resulting in inadequate surface coverage at well centers, and were also observed to delaminate more easily from well surfaces, especially at higher crosslink ratios when pCD tended to contract more upon curing. Following plasma treatment, coatings covered the entire surface of each well, regardless of the HDI crosslink ratio. Similarly, microscope images demonstrated that pCD coatings beaded up and covered very little of the untreated PP suture surface, while they were able to spread and coat a much larger fraction of the surface on plasma treated sutures. This indicates that plasma treatments of PP substrates improve pCD coating uniformity. Taken together with the PP wettability findings, this is likely a result of the enhanced substrate wettability which promotes spreading of the polar pre-polymer mixtures on the surface.

**Figure 4:**
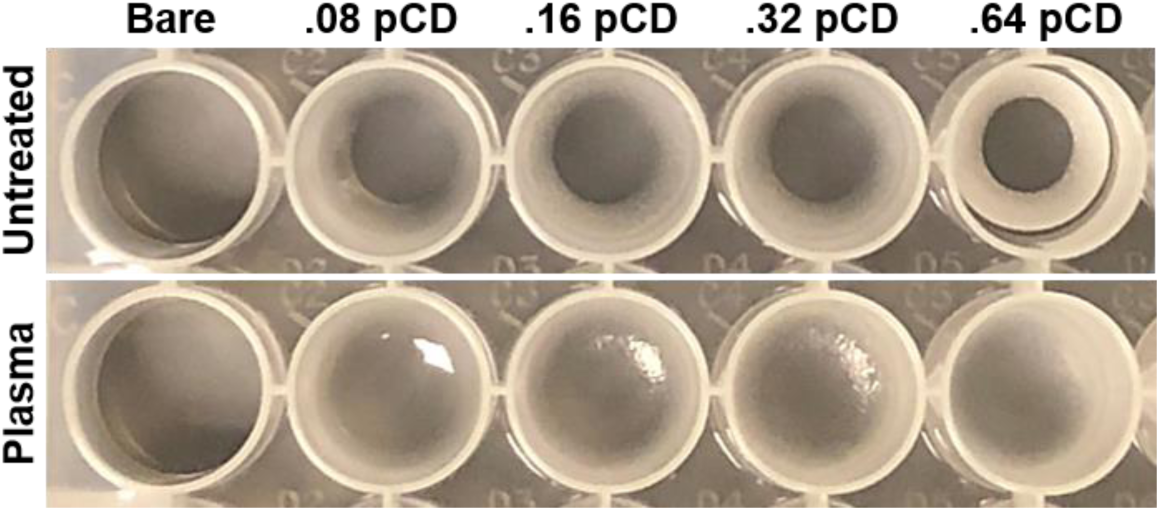
Plasma treatment of PP substrates for 10 min improves macroscopic pCD coating uniformity on wells of a PP 24-well plate. This remained true regardless of the amount of HDI crosslinker added.

**Figure 5:**
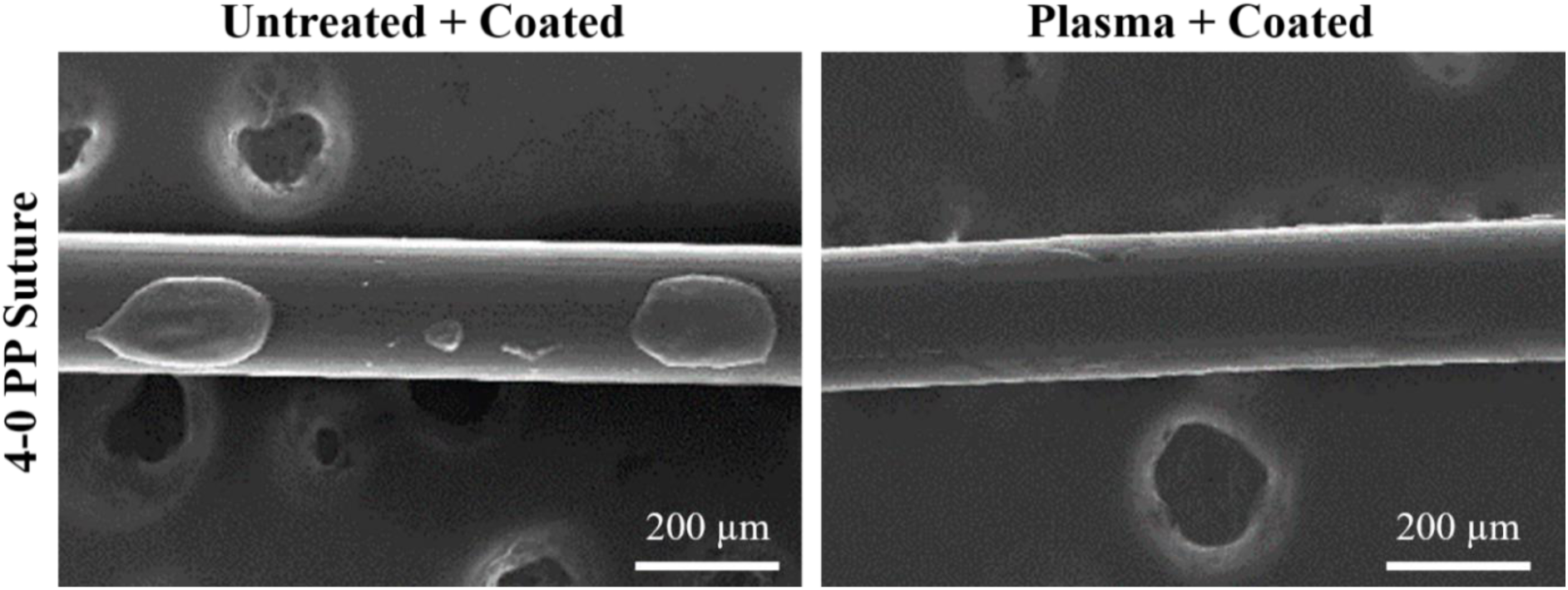
Electron micrographs reveal that plasma treatment of PP suture for 10 min improves microscopic pCD coating uniformity. A thin, uniform pCD coating is apparent on the plasma-treated suture, based on the presence of cracks caused by sample drying upon sputtering and imaging.

Coating adherence was investigated using lap-shear testing, an approach used for measuring the bond shear strength of coatings and adhesives. Lap-shear testing revealed that plasma treatment of PP substrates for 10 min increased pCD coating lap-shear strength by 43% (p < 0.004) (**Fig 6a**) and doubled lap-shear toughness (p < 0.03) (**Fig 6b**). For reference, the lap shear strength of cyanoacrylate-bonded untreated PP adherends is reported to be 0.22 MPa^24^, which is equivalent to the strength observed here for untreated PP bonded with 0.32 pCD. The lap-shear strength and toughness relate to the peak and total area under the load-displacement curve (**Fig 6c**), respectively. The enhanced coating adherence following plasma treatment is considered to result from the evident engraftment of hydroxyls onto the PP surface, which may react to form covalent urethane linkages between the coating and substrate upon isocyanate crosslinking^25,26^. As detailed below, a final experiment was performed to evaluate the validity of this explanation.

**Figure 6:**
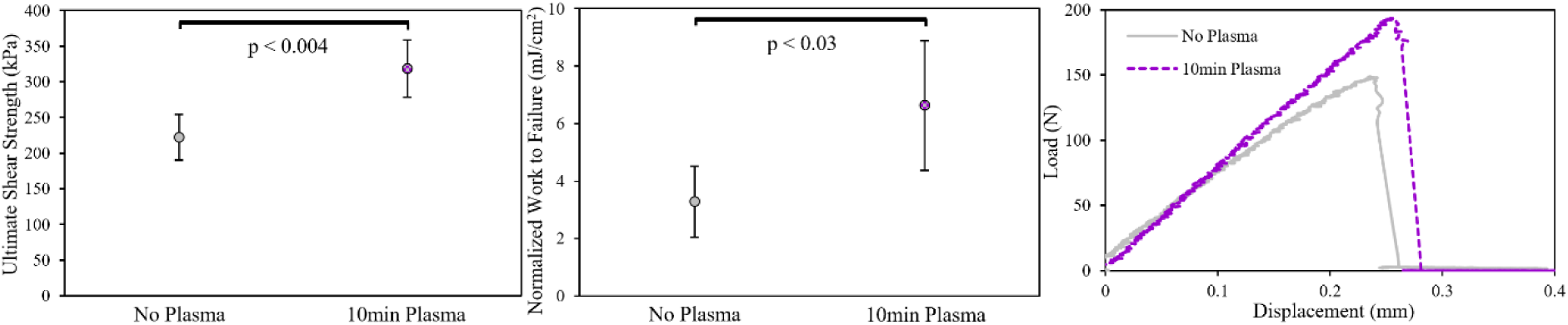
Plasma treatment improves pCD coating adherence to PP substrates in terms of lap-shear strength (a), and toughness (b). Each point represents n = 5 lap joints. Representative (i.e. closest to the average) load-displacement curves demonstrate this trend (c).

### 3.3. Effects of plasma treatment on covalent bonding at coating-substrate interfaces

Having characterized the effects of plasma treatments on pCD coatings, we next aimed to evaluate the formation of covalent urethane bonds between functional groups at the coating-substrate interface. This was done using XPS to detect fluorine as a unique marker of an isocyanate compound, 2-TPI, after exposure of this compound to untreated and plasma-treated PP surfaces followed by copious rinsing. Incubation of untreated PP surfaces with 2-TPI resulted in negligible increase in fluorine content on the surfaces (p = 0.5), while plasma exposure of PP prior to 2-TPI incubation led to a significantly increased fluorine content on the PP surface following incubation (p < 0.005) (**Fig 7, Table 2**). This indicates that plasma treatments of PP substrates enable covalent linkages between the PP surface, most likely via hydroxyl groups, and compounds containing isocyanate groups, such as pCD coatings during HDI crosslinking. This result corroborates the finding that plasma treatment improved the adherence of pCD coatings, and implies that this effect can be attributed, at least in part, to covalent bond formation at the coating-substrate interface. Finally, plasma treatment for 10 min in this experiment resulted in slightly lower oxygen content on the PP surface than in the previous study (**Table 1**), likely as a result of the different rinsing procedure and the increased lapse of time after plasma treatment before XPS was performed.

**Table 2:** Effect of plasma treatment on isocyanate-mediated fluorination of PP surfaces. *Significant difference to non-isocyanate-exposed control.

| Plasma Duration (Minutes) | Isocyanate Exposure | Atomic % Carbon | Atomic % Nitrogen | Atomic % Oxygen | Atomic % Fluorine |
| --- | --- | --- | --- | --- | --- |
| 0 | - | 99.34 ± 0.37 | 0.42 ± 0.02 | 0.25 ± 0.35 | 0.00 ± 0.00 |
| 0 | + | 99.55 ± 0.64 | 0.00 ± 0.00 | 0.15 ± 0.21 | 0.31 ± 0.43 |
| 10 | - | 87.09 ± 0.11 | 0.89 ± 0.06 | 11.97 ± 0.26 | 0.07 ± 0.09 |
| 10 | + | 90.41 ± 0.49 | 0.72 ± 0.32 | 7.48 ± 0.53 | 1.39 ± 0.08* |

**Figure 7:**
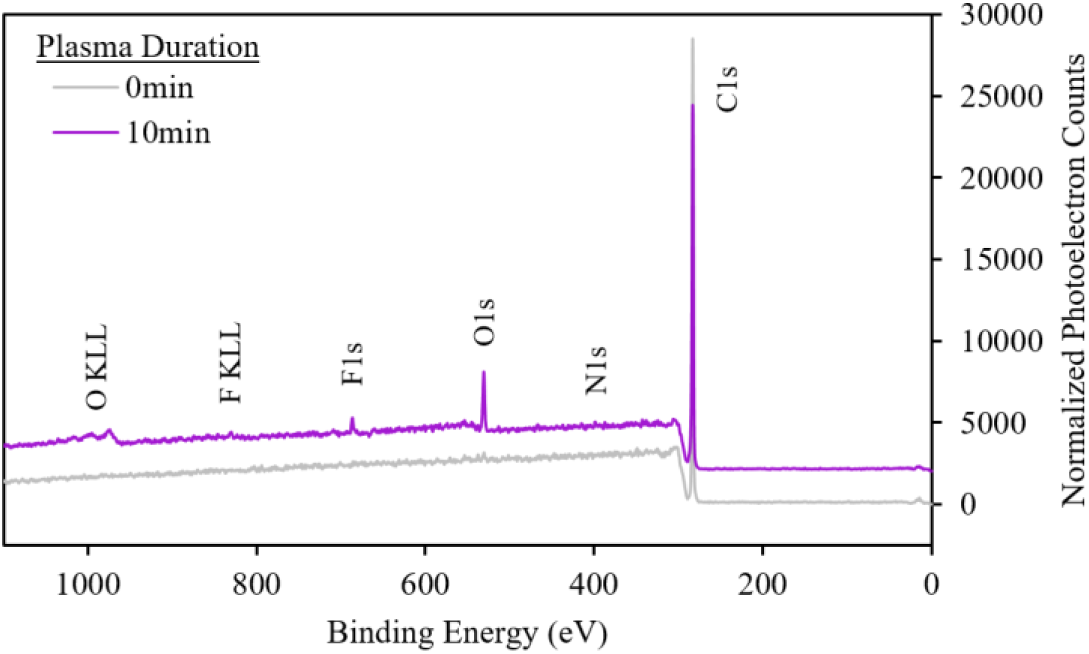
Representative XPS survey scans (rescaled to normalize areas under curves) of PP after 0 or 10 min plasma exposure. Increased isocyanate binding is indicated by the appearance of an F1s peak for the 10 min plasma sample. KLL peaks were not used for analysis. Spectra are intentionally offset along the ordinate.

## 4. Discussion and Conclusions

Uniformity and adherence to the substrate are crucial for the performance and function of any coating. For example, improper adherence of paint to a metal component for a vehicle could accelerate corrosion and dangerous structural failure of the part. Similarly, a lack of adequate substrate coverage of an anti-biofouling coating on a medical device or ship hull could enable organisms to exploit coating defects and attach to the substrate, potentiating problems such as infection or increased drag, respectively. Likewise, delamination of a polymer coating on an intravascular implant could pose risk for lethal thrombosis or embolism^27^.

This study was designed to test the hypothesis that nonthermal plasma treatment enhances the uniformity and adherence of pCD coatings on PP substrates. The findings from contact angle goniometry, XPS, SEM, and lap-shear testing all support this hypothesis. Plasma activation was found here to be a suitable strategy for improving uniformity and adherence of polymer coatings such as pCD, on inert and otherwise incompatible polymer substrates such as PP. This conclusion is in agreement with findings from previous studies that have evaluated effects of plasma on adherence of other polyurethane coating materials to various other low surface energy polymer substrates^26,28^. Martinez et al. demonstrated that atmospheric plasma treatment of acrylonitrile-butadiene-styrene and silicone substrates could improve the adhesion of polyurethane-based paints^28^. Likewise, Bao et al. demonstrated that atmospheric plasma treatment engrafts hydroxyl and amine groups onto styrene-butadiene-styrene rubber surfaces, and that this specific change in surface chemistry, more so than incorporation of chlorine functionalities, enabled increases in T-peel strength of polyurethane coatings after addition of an isocyanate-terminated hardener^26^. Extending the knowledge gained from these prior studies, through detailed XPS-based surface analysis in combination with mechanical lap-shear testing, this investigation has revealed direct evidence for covalent bond formation at the interface between a polyurethane coating and plasma-treated polymer substrate, clarified the basis for these interfacial covalent connections, and identified the contribution of this process to improved adhesion.

The lack of a need for hazardous chemical treatments is an advantageous aspect of substrate plasma activation, especially for coatings to be applied to polymeric biomedical implants, such as PP sutures and hernia meshes. Appropriate selection of plasma treatment parameters can ensure minimal impact on the bulk mechanical properties of such load-bearing substrates^6^. Apart from biomedical purposes, this research has important commercial applications as well. For example, findings indicate that substrate plasma activation could be highly suitable for improving quality of paints or adhesives (especially polyurethane-based formulations) upon low surface energy substrates such as polyolefins or rubber. Likewise, PP surfaces are commonly used in water treatment and food processing despite their high susceptibility to uncontrolled bacterial biofouling^29–32^, while pCD coatings can mitigate bacterial attachment and colonization by way of either active (i.e. antibiotic delivery)^9^ or passive^11^ mechanisms.

Future studies could investigate the effects of plasma treatment on pCD adherence to a wider range of polymer substrate materials, and pCD coating adherence over long periods after plasma exposure. The effects of plasma treatment on the surface chemistry and wettability of a bare substrate are known to diminish (i.e. age) over time. Prior studies suggest that hydrophobic recovery of bare plasma-treated PP normally occurs over the course of several weeks, the rate being dependent on environmental factors and polymer crystallinity^33–36^. However, the application of a covalently anchored coating may limit conformational changes and reorientation of the activated substrate surface, in theory preserving the adherence of the coating long term.

## 5. Acknowledgments

The authors gratefully acknowledge support from National Institutes of Health: NIH R01 GM121477 (HvR), and NIH Ruth L. Kirschstein NRSA T32 AR007505 Training Program in Musculoskeletal Research (GDL). Additional support was provided by the Center for Stem Cell and Regenerative Medicine Student Summer Program (ENGAGE) at Case Western Reserve University (EJL). Core facility services provided by the Swagelok Center for Surface Analysis of Materials and the Advanced Manufacturing and Mechanical Reliability Center at CWRU are also appreciated. The authors also thank the Advincula group for use of the contact angle goniometer, Kevin Abbassi for expertise and assistance with XPS, Kathleen Young for expertise and assistance with SEM, Katherine Yan for assistance with sample preparation, and Erika Cyphert and Nathan Rohner for revision suggestions.

